# The MicroTron: a microfluidic platform for single cell studies in *P. patens*

**DOI:** 10.64898/2026.06.30.735479

**Authors:** Jordi Floriach-Clark, Viola Willemsen

## Abstract

- The effect of some bioactive compounds on living organisms is dependent on their concentration and gradients, as is the case of hormones and signalling peptides, determining cell identity, activity and organism development.
- There are a handful of methods that allow to produce spatially confined peaks of concentration local application of biochemicals on plants, such as agar blocks and microinjection, but they lack in precision, throughput and/or simplicity.
- We developed the MicroTron, a microfluidics-based method specifically for filamentous organisms or life cycle stages, like the moss plant *Physcomitrium patens* protonemata, that serves as a platform for the application of chemicals on single cells and study the cell response.
- We show how chemical applications could be performed on cells, either on the side or apically with dyes and hormones, targeting the cell wall, cell membrane, cytosol and nucleus.
- Treatments could be applied on single filaments and with a precision of up to single cells in optimal conditions.
- This method could be used to study live responses to chemicals with high spatiotemporal resolution.

## Introduction

The organisation of cells in three dimensions to build plant organs requires fine control of the cell division activation and orientation. Such control relies on the cell’s perception of intrinsic and extrinsic spatial information (Smith, 2001; Damme *et al*., 2007). Extrinsic information includes both mechanical and chemical cues, which can originate from the environment or neighbouring cells (de Keijzer *et al*., 2021). Providing those signals to cells exogenously and with high spatial and temporal precision can facilitate the study of the effect chemical cues and their transduction into the cell. This insight could unlock a deeper understanding of the fine regulation of plant cell division control machinery and other processes, as demonstrated in animal cell research (Chen *et al*., 2020; Horowitz *et al*., 2020; Holler *et al*., 2024). However, the translation of animal research methods to plant cell biology is far from trivial. The scarcity of tools with cell precision dedicated to plant systems may be explained by (1) the big size of model plant organs and (2) the stiffness of their cell walls needed to contain the plant cell’s high osmotic pressures, two characteristics that defy the interaction with and targeting of single cells in their natural cellular context. For this reason, we set off to develop novel methods to deliver biochemicals to single plant cells that overcome these limitations to provide a new approach for high precision biochemical application.

For a given biochemical, local and broad treatments can give rise to different outcomes. For instance, local and broad auxin treatment induced different growth fashion (Michniewicz *et al*., 2007). Generally, *treatment locality* consists in the confinement of treatment application to a portion of the totality. At the organ level, locality can be understood as the delivery of a chemical to a group of cells in a specific subregion of the organ; at the cell level, locality refers to the delivery of chemicals to a specific side of the cell. Considering that bioactive chemicals affect all the cells they can reach and induce their effects, such as (in)activation of cellular processes like gene expression (e.g. hormones, peptides, etc.), the locality of the method of application is key to the outcome (Morejohn *et al*., 1987; Etchells & Turner, 2010; Mähönen *et al*., 2014). The diffusion rate and transport mechanisms of the applied biochemical also play a role in the outcome, although these are influenced by biological processes. The combination of treatment locality and compound dynamics can lead to chosen concentration gradients, which have important roles in development as shown with CLE-41 peptides and pH (Scott & Allen, 1999; Etchells & Turner, 2010). The presence of local peak concentrations of nutrients can also induce new organ formations (Remans *et al*., 2006). However, some gradients are short-lived due to rapid transport, such as auxins (Etchells & Turner, 2010).

Some strategies to locally apply chemicals and adjust the cellular microenvironment of intact plant cells existed for long, yet they remain underexplored. The application of local treatments has previously been done at the scale of multiple cells with agarose blocks or arabic gum patches (Ketelaar, 2002; Remans *et al*., 2006; Michniewicz *et al*., 2007). Using a microinjection platform, biochemical applications in and on single cells was achieved for root hairs (Ketelaar *et al*., 2002; Ketelaar, 2002; Wang *et al*., 2015; Baskaran *et al*., 2016). These insightful studies benefited from the precision of the technique, despite their demand for high skill and time investment for a limited throughput. Furthermore, piercing of the cell membrane and wall caused cell arrest, limiting the potential of the test (Hajduk *et al*., 2023). Most of these methods do not provide the simultaneous spatial and temporal control required for valuable high precision studies. In this context, microfluidics initiated a new chapter in the spatiotemporal precision and throughput of (bio)chemical treatments in plant biology, thanks to its versatility of design, high reproducibility, low cost, multiplexing and cell viability improvement for post-treatment long-term imaging (Horowitz *et al*., 2020).

*Arabidopsis thaliana* roots were the first reported plant organs introduced in a microfluidic device, monitored and treated both transversely and longitudinally (Meier *et al*., 2010; Grossmann *et al*., 2011; Stanley *et al*., 2018). Despite the millimetre-scale size of roots, relatively big for common microfluidic devices, the focused-flow transverse hormone treatments were effectively localised in a width down to 10 µm and could be rapidly interrupted or exchanged (Meier *et al*., 2010). On the other hand, the longitudinal flow treatments allowed asymmetrical treatments in each half of the root, providing valuable data of the local response to asymmetric biotic and abiotic cues, which could be exchanged in under 1 minute (Stanley *et al*., 2018; Allan *et al*., 2023, 2024). Pollen tubes were the first cell-scale plant system cultured in microfluidics, and were used to quantitatively study chemotropic growth towards the ovule (Horade *et al*., 2013; Yanagisawa & Higashiyama, 2018).

The moss *Physcomitrium patens* grows in a filamentous fashion, similar to pollen tubes, in the early part of its lifecycle and has received increasing attention in plant developmental biology thanks to its simple body plan (Rensing *et al*., 2020; Floriach-Clark *et al*., 2021). After germination from the spore, the moss grows as single cell-wide filament (i.e. protonema), with widths of circa 20 µm (Floriach-Clark *et al*., 2021). These protonema cells and later stage tissues were successfully cultured, monitored and phenotyped in big chamber microfluidic devices, with successful confocal evaluation of general biochemical treatment applications (Bascom *et al*., 2016). TIRF-like microscopy was proven compatible with slightly compressed protonema in microfluidic devices, coupled with biochemical treatments that could be washed out after the desired time (Kozgunova & Goshima, 2019). These successes in moss microfluidics pave the way to higher spatial treatment resolution, a valuable step towards asymmetric delivery of biochemical cues and cell-to-cell transport studies (de Keijzer *et al*., 2021). Conveniently, *P. patens* protonemata showed the capacity to grow under substantial constriction without effects in microtubule growth speed, and could even adaptively grow inside channels narrower than their natural cell diameter (Yanagisawa *et al*., 2017; Kozgunova & Goshima, 2019). This capacity is valuable for microfluidic platforms which base their functionality in arrangements of channels.

Here we have set out to exploit the filamentous growth of moss to engineer a local chemical delivery system of (bio-)chemicals to single plant cells in their cellular context. Our system takes advantage of the pairing of the strengths of microfluidic devices and the convenient growth fashion and cell characteristics of the plant model system *P. patens*. We demonstrate how this system allows high precision biochemical delivery with cell-scale locality and pre- and post-treatment monitoring at least up to 36 hours. Various locally applied dyes were examined with confocal microscopy to provide visual proof of concept of delivery to cell wall, membrane and cytosol. Furthermore, we demonstrate a biological response to local auxin treatment, proving the system’s capacity to monitor live responses in a biologically relevant example.

## Results

The method of local application we propose is a microfluidic device conceived for organisms that exhibit filamentous developmental stages. Upon emergence from the spore, or when regenerating from homogenized colony tissue (FIG.1A), *Physcomitrium patens* grows as protonemata (FIG.1B), a tip-growing and branching filament that will eventually become an entangled mat and produce the bigger gametophores (FIG.1A,1B). The protonema exist in two types: chloronema and caulonema. The first has cells with an average length of 100 µm, a width of 20 µm, numerous chloroplasts, perpendicular division planes, frequent branching, and a slow growth rate of roughly 2 cells/day or 200 µm/day and serves the function of colony establishment and early photosynthetic capacity. In opposition, caulonemata are a derived exploratory filament present in nitrogen-depleted conditions, that is chloroplast-poor with much thinner and longer cells (down to 10 µm and above 150 µm, respectively), exhibiting higher variability and more extreme cell phenotypes as nutrient starvation prolongs. Their cell division planes are oblique, branching mostly absent and the growth rate is substantially faster. Eventually they may mature to chloroplast-rich phenotypes or branch to make a new transition into (secondary) chloronemal cells. We selected the moss chloronema as an early and lower variability cellular model system to test a microfluidic device approach to deliver (bio)chemicals to single plant cells. Therefore, we optimized the design features to their cellular characteristics.

**FIGURE 1.**
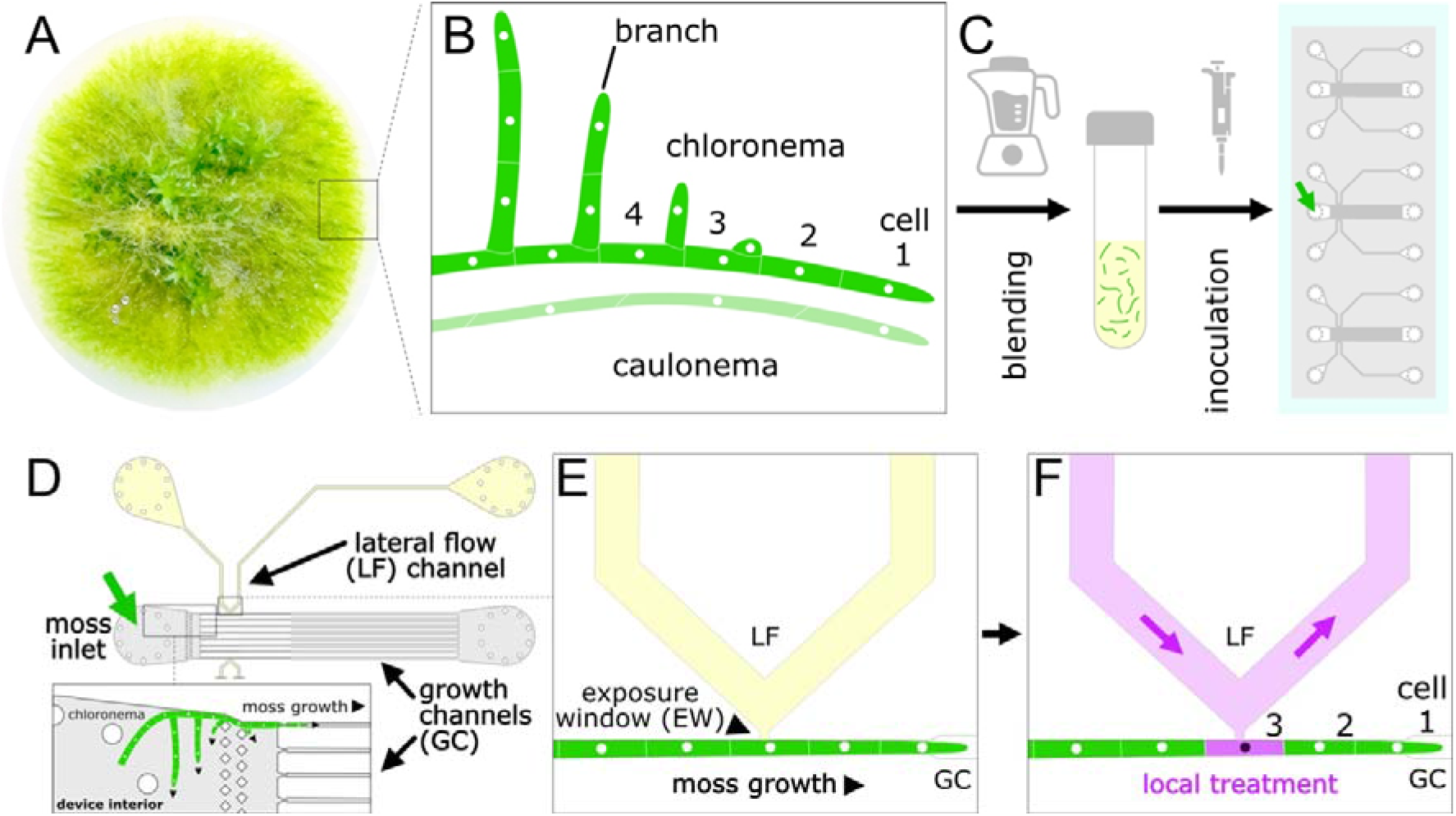
Graphical Abstract of principles and usage workflow of the Lateral Flow microfluidic system with *Physcomitrium patens* (moss) chloronemata. (A) Moss colony showing the mat of protonemata growing outwards. (B) Schematic of chloronema and caulonema types of cells and cell number (1 = apical or tip, 2 = subapical, 3 = sub-subapical, etc.). (C) Process showing the simple chip inoculation process by blending the moss in liquid medium and injecting it into the channel outlet with a pipette. (D) Schematic of the microfluidic device and how chloronema tissue colonizes the chambers and growth channels (black arrow = direction of tip growth). (E) Close-up schematic of the lateral flow-growth channel intersection, with the exposure window connecting both and leaving a single cell exposed. (F) Schematic of the concept of lateral flow-delivered local treatment of single cells.

The proposed design of the microfluidic device allows for the growth of chloronemata from a tissue inlet into eight parallel growth channels (GC), narrow enough to only have room for one filament (FIG.1D, gray). The six inner GC are isolated along their whole length, and the outer two GC intersect tangentially with two lateral flow (LF, FIG.1D, yellow) channels through which a biochemical solution can be delivered. This intersection consists of a V-junction connected by a small opening or exposure window (EW) that provides cellular surface access to the growing protonema (FIG.1E). This design provides 2 GC for treatment and 6 GC as control per experimental unit (FIG.1D). The design dimensions of the EW (LF-to-GC junction) and GC width and height are specifically selected for cells to seal off the EW with their natural cell turgor. The subsequent isolation of the LF channel allows the treatment to flow through to the outlet and treat the cell located at the EW individually (FIG.1F). Making the EW smaller than the cell length can theoretically minimise the total treated surface down to subcellular scale, thus increasing spatial precision or treatment locality.

The high regeneration capacity of moss allows to homogenise the moss tissue and directly inoculate it into the moss inlet of a microfluidic device prefilled with liquid medium (FIG.1C, 2A) as reported previously (Bascom *et al*., 2016; Yoshida & Kozgunova, 2023). After 48 hours in standard culture conditions, the moss will initiate new cell divisions and protonema will colonize the channels, including the EW test area (FIG.2E-2H). Media refreshment is not necessary, only a saturated environment to prevent drying (FIG.2B-D, 2K, Supplementary FIG.1).

**FIGURE 2.**
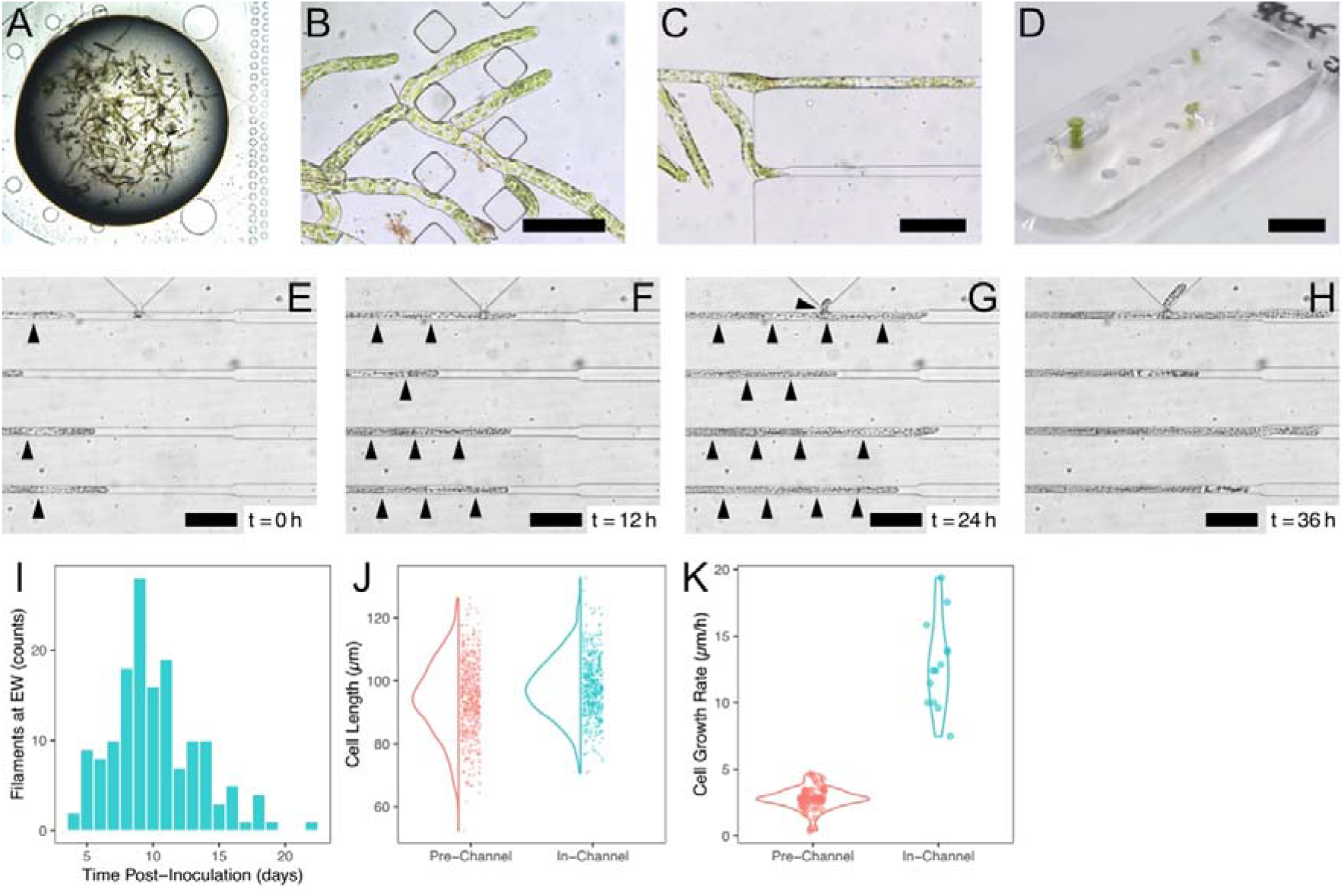
Growth and performance of *P. patens* in our microfluidic device design. (A) Chip inoculated at t = 0, showing green blended chloronemata in the inlet. (B) Growth of regenerated chloronemata 48h after blending. The filaments start colonizing the space in the pre-channel chamber. (C) Colonization of growth channels. (D) Appearance of the device after a 10 dpi. (D-H) Time-series of chloronemata growing in parallel GC. Black arrows indicate cell divisions. (I) Frequency plot of time needed for moss to reach the EW after chip inoculation (n=152). (J) Cell length distribution in chips before (n=532) and after entering the channels (n=465). (K) Growth rates of before (n=58) and after (n=12) entering the channels. Scale bars = 50 µm.

The time for chloronemata to enter the channels depends on where the blended cells landed upon chip inoculation. Growth will initiate in the open chambers preceding the channels (FIG.2B). The moss grows in all directions without apparent preferential direction. Within 9±3 days after inoculation 70% of subapical chloronemata cells in the outer LF-GC reach the EW, raising to 90% after 20 days post-inoculation (FIG.2I). The remaining 10% EWs are spatially inaccessible due to obstruction from other filament given that the device height (19.2 µm, Suppl. FIG.2, Suppl. Table 1) only allows room to a single layer of chloronema cells.

The chloronema cell characteristics, such as cell length, within the chamber prior to the growth channels are largely unaffected. As reported in previous literature, a chamber height of 8.5 µm had no effect in microtubule polymerization velocity, and our device has a height is well above it (FIG.2J, Suppl. FIG.2., Suppl. Table 1; (Kozgunova & Goshima, 2019; Floriach-Clark *et al*., 2021)). However, the growth inside the GC could be affected by the mildly constrictive characteristics both in heigh and width, and might induce deformation, stress or growth defects. For instance, we report the sustained inhibition of branching inside GCs (FIG.2E-H). Indeed, our measurements show that the cells have very similar cell length inside and outside the channels, with only minor but significant differences of cell length outside (95±11 µm, n=532) and inside (98±9 µm, n=465) channels (Welch Two Samples T-test, p = 2·10^−7^; FIG.2J). Growth rate analysis further confirms that cells remain robust and actively growing under confinement, with values on the same order of magnitude as previously reported for chloronemata (5.4–7.2 µm h-^1^) and caulonemata (16–20 µm h-^1^ (Bascom et al., 2016; Rawat et al., 2017; Bibeau et al., 2021). Specifically, cells outside (2.8 ± 0.8 µm h-^1^, n = 58) and inside (12.8 ± 3.3 µm h-^1^, n = 13) channels both display sustained growth (Fig. 2K). We did not aim to systematically quantify the growth rate differences in multiple conditions. We interpret that the degree of constriction in the channel is biologically innocuous to the normal development of the main filament cells, as our designed aimed to achieve.

As cells grow along the outer GCs, we observe that the release of growth constriction at the EW promotes branching events, with a frequency of 47% branching in (n=34, FIG.3A) and 24% not branching (n=17, FIG.3B), and a quarter of cases showing filaments turn into the LF channel or channels remain empty (n=21; FIG.3C). This variability provides two cellular states to which treatments could be applied: a branching cell (apical treatment) and unbranched cell (side treatment). These positions are biologically distinct: the cell walls of the tip-growing branch cells are developing, and membranes are formed, leading to a different cell wall, and membrane composition (Bibeau *et al*., 2021). In addition, these cells have highly active endocytic/exocytic pathways and show high metabolic rates. On the other hand, unbranched cells are less metabolically active and have a fully developed cell wall (Supplementary FIG.4). Thus, we regard apical and side applications as biologically distinct cases which may serve as unique testing conditions.

To test the precision of the device we first tested the application of the cell wall dye Calcofluor White (CFW), which binds to crystalline cellulose microfibrils (Bidhendi *et al*., 2020). As shown in FIG.3E F, the application occurs mainly within the EW, in both apical and side applications, and occasionally beyond it in side applications (FIG.3D). The treatment locality varies from 0 (single-cellular) to ±1, ±2 or ±3 cells (multicellular), in all cases staining only the cells neighbouring the EW. The result variability may have device characteristics and biological origin. We measured that our devices’ channel dimensions are close to those by design and very regular, showing a width of 16.5±0.2 µm and height 19.0±0.2 µm. We also evaluated the cross-section profile, which showed wall bending and increased width at the contact with the glass bottom (Supplementary FIG.2, Suppl. Table 1). These wider corners at the base may explain the occasional CFW escapes by diffusion through the corner of the channel, which could vary from batch to batch with differences in adhesion, and prevent that the protonemata blocks the EW completely. Apical application of CFW showed how locality can be single-cellular possibly aided by the additional blocking of the cell growing into the treatment channel (FIG.3E).

**FIGURE 3.**
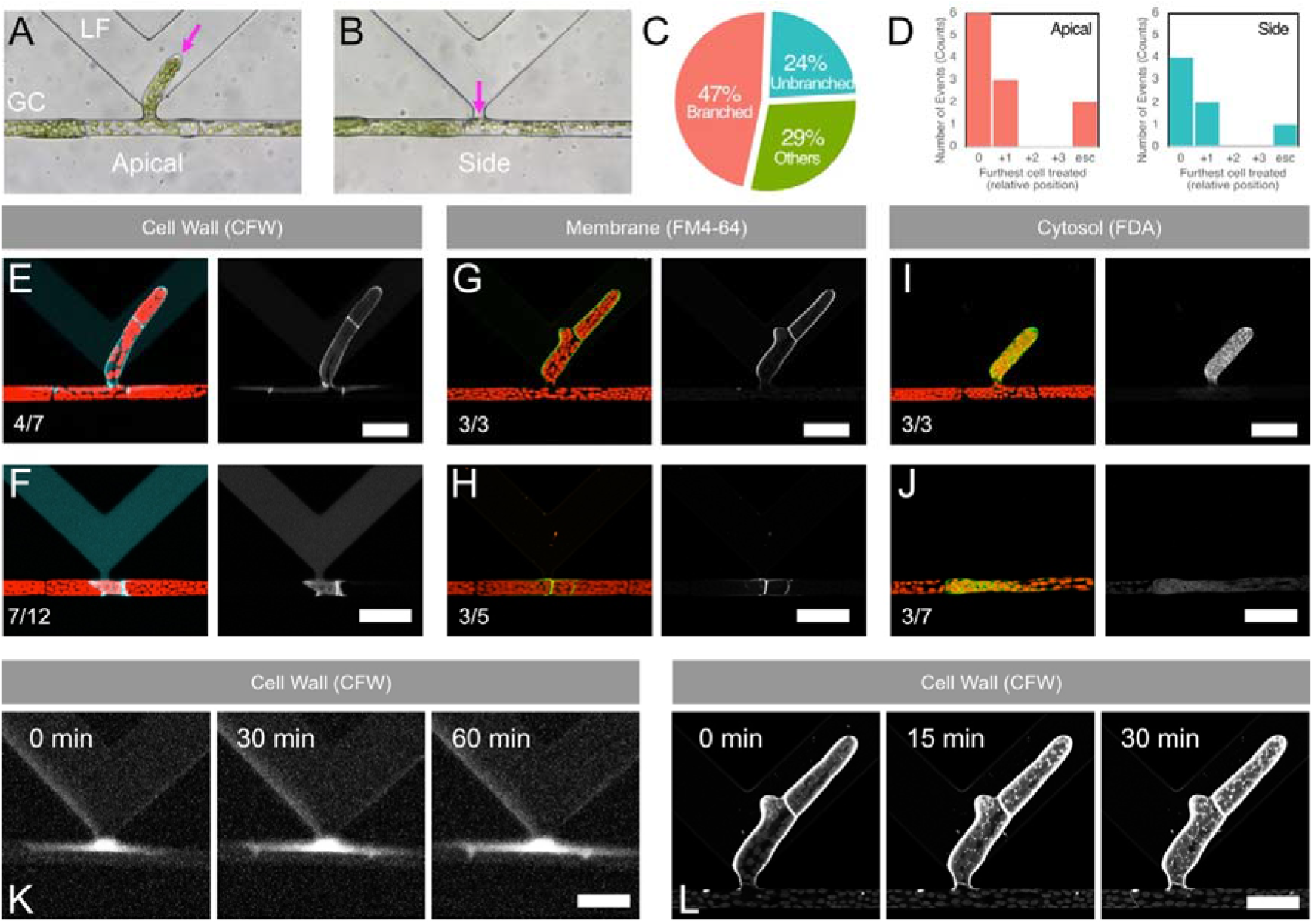
Chloronemata individual cells could be stained, both at apical and side locations. (A) Chloronema in GC and branching into the LF channel. (B) Chloronema in GC that does not branch in. The pink arrows indicate main exposure points. (C) Summary of probability of branching (n=72). (D) Locality of the applied treatment, where 0 represents single-cell treatments, ±n refers to cells treated beyond cell at EW, and ‘esc’ means escape for apical (n=11) and side (n=7) applications. (E-L) Confocal images of cells locally treated with several dyes and cell states. Scale bars = 50 µm. (E-F) Calcofluor White (CFW) chloronema cell wall stains, to apical (E) and side (F). (G-H) FM4-64 chloronema membrane stains, to apical (G) and side (H). (I-J) Fluorescein Diacetate (FDA) chloronema cytosol stains, to apical (G) and side (H). (K) Time-series of CFW local stain showing treatment diffusion. (L) Time-series of FM4-64 apical local stain showing dye internalisation into endosomal system.

**FIGURE 4.**
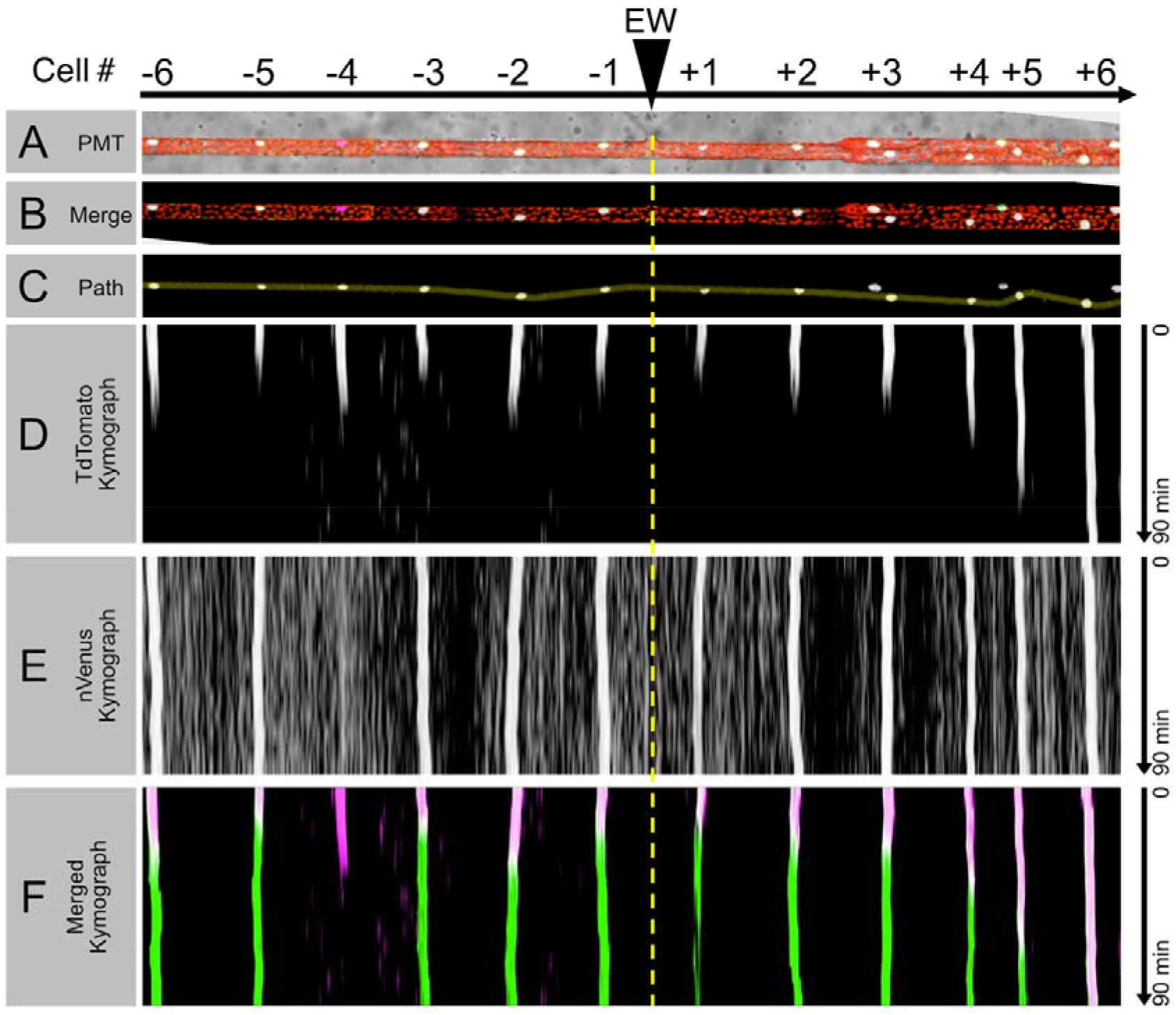
Application of auxin on an auxin-sensing R2D2 reporter line, a bioactive hormone, inducing local and global biological response. (A) Merge of PMT and all fluorescent channels of the exposed filament at t = 0. (B) Merge without PMT. (C) nVenus channel with the trace of a path for the later kymogram. (D) nTdTomato channel kymogram, showing a decrease of signal over the course of the following 30 minutes. The sooner the auxin response occurs, the shorter the signal decay. (E) nVenus channel kymogram showing no nuclear signal decay as control. (F) Merge of nTdTomato and nVenus.

Treatment locality was further confirmed with the membrane stain FM4-64 and cytosolic vitality stain FDA. In apical applications, exposed cell membrane and cytosol were immediately and locally stained at single-cell resolution, respectively (FIG.3G,3I). For FDA, a signal builds up upon hydrolysis by intercellular esterases. We detected signal over time in the apical application, and beyond it with minimal intensity. For side application of both FM4-64 and FDA, we observed low signal intensity at the treated cell, and higher concentrations were required to obtain high signal intensity (FIG.3H,3J). When replicating this treatment, some cells did not show the expected staining. Subsequently, we checked whether all cells respond equally with the bulk application of dye and confirmed this is not the case, especially with FM4-64 and FDA, which can vary dramatically from cell to cell (Suppl. FIG.4). In FIG.3D, a frequency distribution is shown of how local the treatments are using this microfluidic-moss system. Most events occur in the directly exposed cell and the cells around it.

The local applications were followed in time and showed how diffusion of dye also plays a role if the dye or treatment is left in the LF channel. In FIG.3K, CFW side application expanded its signal over the subsequent hour, whereas FIG.3L shows how FM4-64 is internalised into the endomembranous system over the following 30 minutes, demonstrating dynamics and the viability of the cell during the observation and treatment. Taken together, these results prove that (bio)chemical treatments can be achieved locally down to single cells and monitored over time to assess effects of high locality treatments.

The capture of cell response to local bioactive chemical treatments such as hormones is particularly valuable. However, the time for the cell to react and produce a detectable change of signal sets hormone treatments apart from the local application of dyes. Therefore, we tested the local application of the auxin indole-3-acetic acid (IAA), a hormone involved in many developmental processes, and in the maintenance of specific cell identities in plants. IAA is perceived in the cell nucleus by Aux/IAA proteins through their DII domain, which form a gene expression inhibition complex with ARF proteins in the absence of IAA. When IAA is present, this complex is rapidly disrupted due to Aux/IAA degradation, and ARF activates transcription of key genes in minutes. The R2D2 reporter system was engineered for *A. thaliana* to exploit this mechanism to rapidly and quantitatively detect the presence of auxin (Liao *et al*., 2015). In *P. patens*, the system consists of two fluorescent reporters (nVenus and nTdTomato), expressed under the same promoter, which have a different DII domain version, one IAA-sensitive (wild-type, DII) and one IAA-insensitive (mutated, mDII). The level of expression of both proteins is the same by use of the same promoter and genomic region, but the final abundance of the IAA-sensitive fusion nTdTomato-DII is conditioned to IAA levels whereas nVenus-mDII is unaffected (Brunoud *et al*., 2012). Upon an increase in IAA concentration, nTdTomato-DII is immediately marked for degradation and a decrease of the ratio nTdTomato/nVenus occurs within 30 minutes of application of IAA (Thelander *et al*., 2019).

We show that side application of IAA induces nTdTomato signal depletion detectable from 30 minutes after hormone application (FIG.4D-4F), while nVenus signal levels stay high and constant (FIG.4E). The treatment was monitored in time for up to 90 minutes, and we saw how the local application of IAA induces a fast decrease of nTdTomato signal in directly exposed cells, with full nTdTomato depletion within half an hour. Interestingly, the kymogram in FIG.4D shows this happening beyond the directly exposed cells. The neighbouring cells in both directions show a decrease of nTdTomato signal that becomes undetectable later than the directly treated cells. This demonstrates how cell-independent dynamic processes can be followed in time and aligned with the known transport of IAA from cell to cell. Branched out chloronemata directly connected to the treated cell in the parallel GC also responded to the IAA treatment, implying transport, whereas control of distant protonemata in other growth channels within the same microfluidic device showed no decrease in TdTomato signal within the same time. However, 24 h post-treatment, all cells in the channel responded to IAA.

## Discussion

We adapted the features of a microfluidic device to *P. patens* filamentous growth fashion and cell shape plasticity. Two days after blending and inoculation of the chips, the chloronemata regenerate as normal and colonize the growth channels. The location where each chloronema cell establishes is permanent, given that cells do not elongate after their formative division, and therefore the filament position is unchanged over time. This characteristic is critical for the compatibility with the proposed device concept. Our design arrangement and size allow two technical replicates per experimental unit, with three of these fitting in a single big coverslip of 24 × 60 mm, similar to standard microscopy slides, for biological replicates. Furthermore, this approach renders the use of advanced equipment such as pumps and control unnecessary (although possible), as it would be necessary for more complex focused flow approach, making it more accessible and easier to leverage by any cell biology laboratory with limited or no previous experience in microfluidics.

Once within channels, side branching was suppressed, but it resumed in new cells upon release of spatial constraints, implying inhibition of branching is limited to constricted cells. Occasionally, outward bending of the PDMS channel walls by the moss cells was observed, which hints to a filling up of all the space available in the channel and some level of flexibility of the channel structure to accommodate cells, and suggests that the spatial limitation has mild effects that the cell mechanics can handle. Other cell parameters like the cell length showed regularity and similarity outside and within channels. Interestingly, the growth rate of chloronemata in the device is comparable to the previously reported in microfluidic devices. Taken together, these data indicate that gentle spatial constriction is perceived but not detrimental to growth. Considering that in natural situations the protonemata can grow through tight slits within the soil or rocks as part of its exploratory and colonizing function, we argue the gently restrained growth occurring in our device is a physiologically relevant and natural growth condition facilitated by its growth plasticity. Thereby, we propose the spatially-challenged growth may represent a more natural situation than standard agar or liquid culture, which may explain the ease of culture of the moss within these devices. Beyond the mentioned parameters, we do not detect signs of cell stress during the first 15 days of culture. From there, some chloronema cells occasionally divide in half or die, a stress-related response that can be attributed to the lack of media renewal, subsequent nutrient depletion and waste product excretion and self-poisoning, although many others stay alive and grow.

After an average of 9 days post-inoculation, the chloronema has grown past the exposure window of the device and apical and side local applications are possible. This was achieved for both apical and side staining of cell wall, membrane and cytosol, and side treatments for nuclear sensing by R2D2. The treatments with FM4-64, FDA and IAA were dynamically observed for 30 to 90 minutes after treatment, showing the living state, activity and responsivity of the cells in these conditions.

The best performing side applications were on third cells (i.e. sub-subapical) or older. From the sub-subapical cells onwards, treatment leakage or escape through the growth channel was observed in a minor fraction of the events, with decreased frequency on older cell applications. Single-cellular treatment applications on the apical and subapical cells of the main filament were not achieved, as escapes occurred in all attempts. We could observe free channel space around the thinner apical cells, whereas subapical cells were wide. When the treatment escapes through the growth channel, several cells are exposed to it, reducing the locality of treatment, and mostly in the apical side (thus, hardly any flow towards older cells).

Given these observations, we speculate about two possible mechanisms by which escapes occur. Firstly, in cases of high spatial constriction and tight cell fitting in the channel where leakage is improbable, this may be a consequence of peak high pressures when treatment is manually and quickly applied using a syringe. Handheld syringe application is convenient and fast, but variability arises from operator and possible obstructions from biological debris, facilitating pressure shocks, which could potentially and reversibly deform the exposed cells or PDMS and lead to wider treatment range on some occasions, even if no escapes occur. Syringe-pumps and similar systems provide more control in these cases by facilitating lower and steadier flowrates. Secondly, biological and channel size variability could cause the growth channels to be incompletely filled, and the small gaps in the corners between the circular cell profile and square channel profile could provide a passage for treatment escape by not locking the possibility of flow (Suppl. FIG.2). As seen by our results, the treatment escaped for side applications on first and second cells of the main filament. Given that protonema exhibit tip growth and no diffuse or secondary cell growth, the first cell may have not reached its final diameter and escapes are expected, whereas second cells must have a mature cell width (FIG.1B,1F; Menand *et al*., 2007). Therefore, from the second cell (FIG.1F) the channel wall and cell wall resistance should be sufficient to stop flow, but due to resistance increase as a function of length, older cells may have less escapes. Following this flow-locking rationale, the apical applications (FIG.3A,3E,3G,3I) should be the ones exhibiting least escapes and highest locality (i.e. smallest treated area), because the cell entering the channel obstructs maximally any flow towards the GC. Conveniently, this event covers the biological case of apical treatments that were not possible in the main filament due to escapes. Due to the possibility of escapes by experimental variability, and for validation of treatment locality, we consider recommendable to co-apply or sequentially apply a dye after any biochemical treatment to verify the profile of exposure in a few replicates.

We noticed that side applications occasionally needed higher treatment concentrations than apical applications to provide high signal. While this may partially be a consequence of treatment adsorption into the PDMS (Wang *et al*., 2012), it may also be influenced by the flowrate profile in the lateral flow channel. Saadat *et al*. (2024) showed that the angle acuteness of channel geometry, such as that of our lateral channel pre- to post-exposure window, affects the flow rate profile particularly at the turn. This may influence the amount of treatment that would reach the cell at the exposure window. The geometry we propose has the most favourable treatment distribution profile thanks to low angle acuteness (90º), in which the flowrate maximum is closer to the exposure window compared to every other geometry simulated. Nonetheless, when considering a single closed channel (with no exposure window to another channel), the flowrate is near 0 at the channel sides. Despite we worked on the assumption that a fully developed chloronemal cell locks the exposure window, a minimal space in the corners of the channel may allow some net flow towards the exposure window, especially at higher pressures. In such case, the successful treatment delivery we observe with our geometry would require higher pressures, higher concentrations and/or diffusion time to reach the cell at the exposure window. Additionally, in static conditions, the small length between the growth channel and lateral channel would allow for diffusion to increase the biochemical delivery over time. Underexposure to treatment was never detected in apical applications in which the cell is exposed directly to the maximum flow rate location. Interestingly, we noticed, especially in the case of the hydrophobic dye FM4-64, that the exposed membrane surface was not homogeneous, showing “brighter” sides (higher signal intensity) at the side of the cell exposed directly to the flow inlet and a “shadow” (low signal intensity) of treatment in the side of the cell towards the flow outlet direction. We envision this may be exploited for asymmetrical treatments in the longitudinal direction within the cell of interest.

The application of IAA gave a local effect and a distant effect as time passed. This may be explained by active transport of auxin by PIN proteins, a well described process in plants. For moss, it has been reported to not have a preferential direction, so all cells of the protonemata should lose nTdTomato signal over time. Interestingly, we could follow the levels of auxin in each cell over time after the local application, and we saw nTdTomato being degraded at different speeds.

In sum, these results prove the live response to a biologically relevant molecule and the response of further cells over time, highlight the potential of application of this device to study compound dynamics, signalling across long distances and cell-to-cell transport. We reckon this microfluidic system could provide a tool to trigger processes where this level of cellular resolution is valuable to answer further biological questions or test (bio)chemical-plant cell interactions with high precision in cell response characterisation.

We achieved to study the development of a plant cell with actionable single-cell resolution in living conditions. Treatment effects were monitored live for hours and can be tracked for days with the right microscopy-culture setup. Treatments were delivered on single cells with immediate effects on them, and later effects on neighbouring cells by means of cell-to-cell transport of biochemicals. This approach paves the way towards precise external cue modification in a plant cell biology model system. Despite it being limited to filamentous organisms like *P. patens*, the gain in precision could allow to unlock access to information that was not achieved before.

## Methods

### Design and Production of Microfluidic Devices

The device was designed using AutoCAD 2023 (Autodesk, California) and adjusted for exposure with KLayout v0.25 (Free Silicon Foundation). Silicon wafers of diameter 76.2 mm (Si-Mat, Germany) were coated with a 20 µm-thick layer of Kayaku SU-8 25 permanent negative epoxy photoresist following manufacturer’s instructions using a Laurell WS spincoater. Resin exposure was done using a MicroWriter ML® 3 Pro (Durham Magneto Optics, United Kingdom) at an energy of 200 mJ/cm^2^ and no global focus correction. Post-Exposure Bake step was modified to 90s@65ºC and 270s@96ºC, cooled down to RT for 120 min, and developed by orbital shaking in pure PGMEA (Sigma-Aldrich, US) for 20 min at 100 rpm, then washing with isopropanol and MilliQ® water, and thorough drying with N_2_. PDMS chips were casted using DOW SYLGARD 184 and baked at 70ºC overnight. Holes were carved using 1-mm wide biopsy punches (KAI, Japan). Chips were plasma-bonded to 60 × 24 mm CORNING nº 1 ½ coverslips as shown in REF. The real dimension of the chip features were verified by filling the channels with a concentrated basic fluorescein solution and confocal microscopy. The measured heigh of the chip measured through this method was 17±1 µm, and the exposure window width (EW) was 12.3±0.7 µm at its widest section.

The devices were sterilised by autoclaving (20 min @ 121ºC, 15 psi) and oven-dried (3 h @ 130ºC). Then they were exposed to 30 minutes of UV light before inoculation.

### Moss culture

#### Genotypes and Propagation

*Physcomitrium patens* wild-type ecotype Reute (Hiss *et al*., 2017) and Gransden (Ashton & Cove, 1977) were used according to experimental requirements. The reporter line R2D2 was produced by Thelander *et al*., 2019 in the Reute genetic background, whereas all other reporter or mutant lines were produced in a Gransden background. Routine propagation was done weekly by tissue homogenization in sterile Milli-Q® water and pouring on cellophane-covered BCDAT or BCD petri dishes, and cultured at 25ºC under continuous white light (Nishiyama *et al*., 2000).

#### Microfluidic device culture

The microfluidic devices are prefilled with liquid BCDAT using a manifold connecting a syringe to the three device inlets for complete filling and gas removal. The tissue cultured for propagation is collected after 5 days into liquid BCDAT and homogenised as a small-concentration tissue suspension. Subsequently, 300 µl of well-mixed tissue blend is introduced in the prefilled devices by the central inlet using a 1-ml micropipette, and pushed until a droplet forms at the central outlet. Excess droplets or other wettings are dried up with sterile tissue paper. The device is secured inside a petri dish with sterile tape by the coverslip and a pool of liquid BCDAT is added around it to keep humidity saturation within the dish. The petri dish is sealed with micropore tape (3M, USA). This is cultured at 25ºC under continuous light for up to 20 days without any media flow needed, with daily monitoring of the growth until the moss protonema reaches the exposure window to the lateral flow. The live imaging for up to 72h was done under a different growth regime, with same continuous light but different spectrum and intensity, and adding liquid BCDAT reservoirs in all inlets and outlets.

### Microscopy

#### Light microscopy

Daily observation was done using a Nikon Diaphot 200 Hoffman Modulation Contrast inverted microscope with light source PSM-2120 and camera Nikon Digital Sight DS-Fi1-U2 (1920 × 1200 px) with NIS Elements 4.60.

##### Confocal microscopy

Observation of local applications was performed with a laser scanning confocal microscope using the Leica Stellaris 5+ inverted confocal microscope equipped with a white light laser (485-685 nm) and solid state lasers of 405, 448, 488, 561 nm which were selected as needed depending on the prove being tested. Leica Application Suite X (version 4.8) software was used both for acquisition and image processing. Live imaging for 72 h is done using an spinning disk confocal microscope Andor Revolution (2013) which has a temperature and humidity control module INU by TOKAI HIT.

### Chemicals

Fluorescent Brightener 28 or CalcoFluor White (“CFW”, CAS 4193-55-9) was purchased from Sigma Aldrich as a 25% m/v water solution and diluted in BCDAT media to 2 mg/ml. FM™□4-64 (CAS 162112-35-8) was purchased from Invitrogen (Thermo Fisher Scientific, Massachusetts) as powder and dissolved to make a 10 µM solution in BCDAT. Fluorescein Diacetate (“FDA”, CAS 596-09-8) was purchased from Sigma Aldrich as powder and dissolved to 5 mg/ml acetone stock solution, then to 20 µg/ml in BCDAT. Indole-3-acetic acid (IAA, 87-51-4) for plant cell culture was purchased from Sigma Aldrich as powder and dissolved at high pH and dissolved in BCDAT to 1 mM stock.

### Biochemical delivery

#### Application

Treatment solution stocks were diluted in BCDAT to the relevant concentration to minimize osmolar stress. Applications were done manually through PTFE tubing 1/16” OD X 1/32” ID microfluidic connected to the selected treatment inlet or outlet (depending on desired direction of application) of the experimental unit to a 1-ml syringe containing the treatment solution and manually applying pressure. In optimal conditions, the treatment circulates with minimal resistance to the treatment outlet, forming a droplet as liquid comes out. The excess droplet is dried with a piece of paper tissue, and treatment is further circulated until five or more droplets have come out and been dried to account for the internal volume of the LF channel, which is approximately of 100 µl. For time-limited treatment exposure, BCDAT is circulated after the selected incubation time using the same method. The device is imaged before and after treatment circulation as control. When possible, dyes that did not overlap in emission spectrum were applied one after the other, and no signal interference was observed.

| Treatment | Abbr. | Target | Composition | Exc. Laser (nm) | Acquisition Window (nm) | Supplier |
| --- | --- | --- | --- | --- | --- | --- |
| Calcofluor White | CFW | Cell wall | 2 mg/ml | 405 | 415-440 | Sigma-Aldrich |
| FM4-64® | FM4-64 | Membranes | 10 μM | 505 | 576-640 | Thermo Fisher Scientific |
| Fluorescein Diacetate | FDA | Cytosol | 20 μg/ml | 495 | 518-545 | Sigma-Aldrich |
| Indole-3-acetic acid | IAA | Cytosol | 1 mM | 514 (mVenus)<br>554 (TdTomato) | 519-553 (mVenus)<br>564-618 (TdTomato) | Sigma-Aldrich |
| Chlorophyll | - | - | - | 495 | 680-770 | - |

### Data analysis

Image analysis was performed with ImageJ (version 1.54g, National Institute of Health, USA) and data processing, statistics and plotting was done using RStudio IDE (version 2025.9.1.401, Posit Software, Boston, USA).

## Acknowledgements

We would like to thank Martijn van Galen and Joris Sprakel for the valuable discussions on the initial design and production of the first microfluidic device. We would like to thank Ben Scheres for the valuable discussions regarding the project and manuscript, and Norbert de Ruijter for his technical advice on microscopy.

## Author Contributions

**Jordi Floriach-Clark**: device design, production, characterization and troubleshooting; device use protocol development and implementation; experimental design and execution, data processing, manuscript writing. **Viola Willemsen**: device design troubleshooting; device use protocol development; experimental design, manuscript reviewing

## Data availability

Under request.

## Supporting Information

**Supplementary Figure 1.**
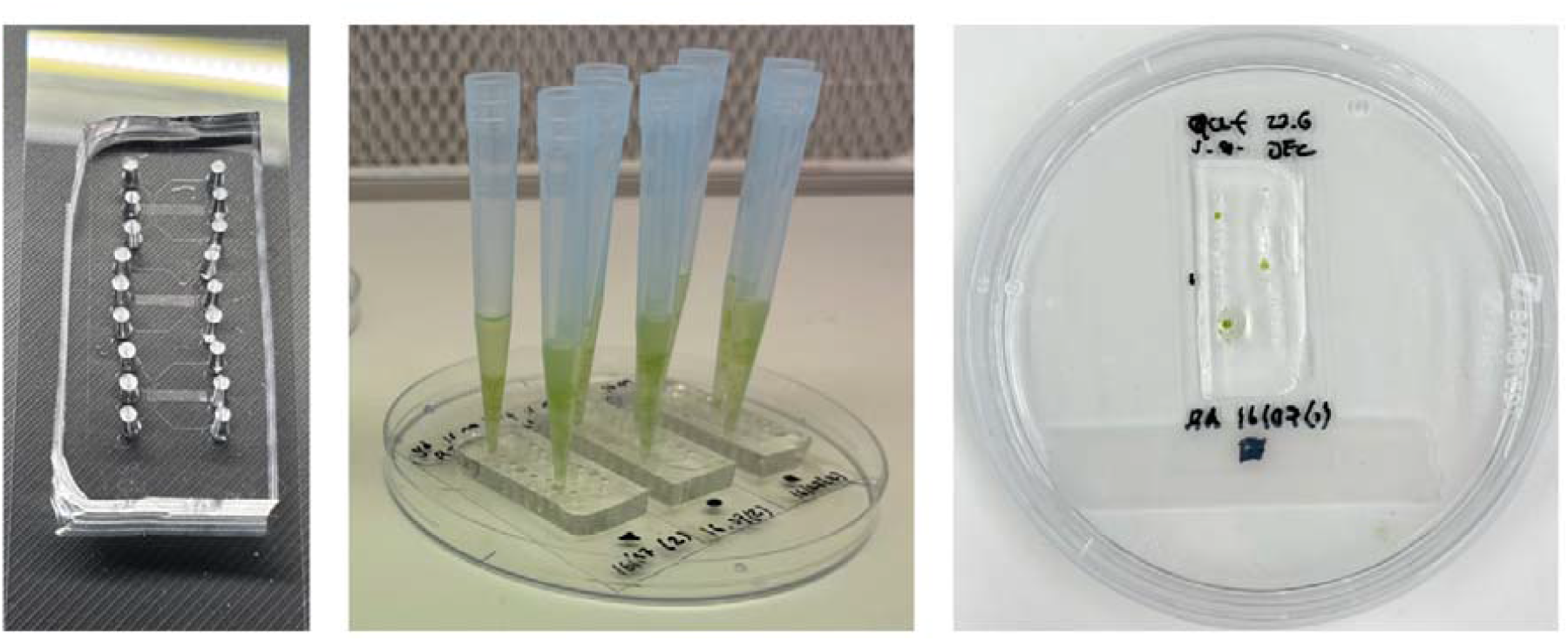
Appearance of the microfluidic device along its use. Device before use showing the grouping of three experimental units (left). Inoculation of moss inlets with blended tissue. Most volume is not used. (center) The device is cultured semi-submerged in media, with the upper surface exposed to air (right).

**Supplementary Figure 2.**
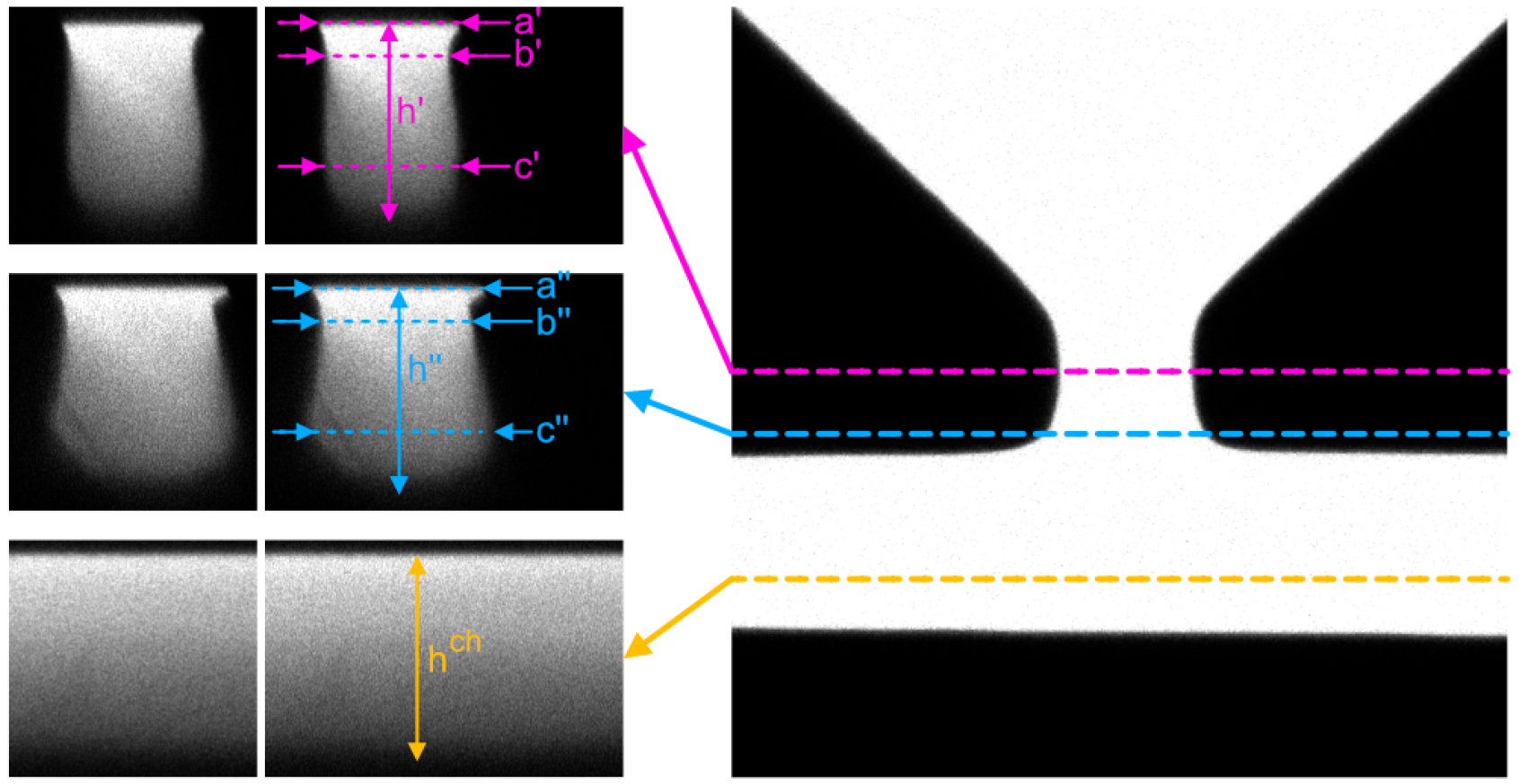
Real channel profile dimensions measured by confocal microscopy of a fluorescein-filled device.

**Supplementary Table 1.** Measured dimensions of device exposure window and adjacent growth channel, in *µm*.

|  | ' | " | ch |
| --- | --- | --- | --- |
| a | 12.7 | 15.6 | - |
| b | 10.8 | 13.4 | - |
| c | 12.4 | 17.3 | - |
| h | 19.2 | 18.8 | 19.2 |

**Supplementary Figure 3.**
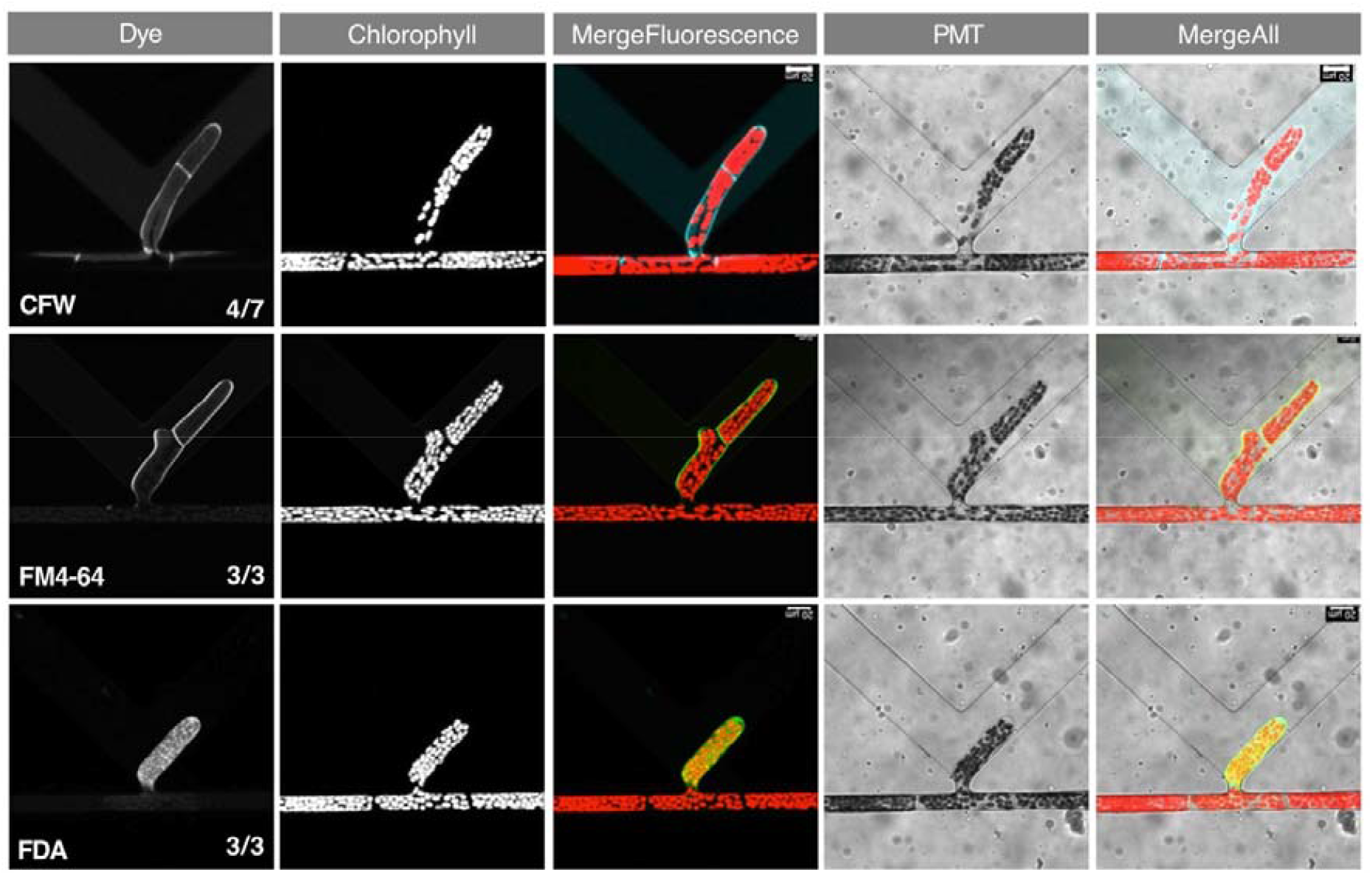
Apical applications of dyes, with all channels.

**Supplementary Figure 4.**
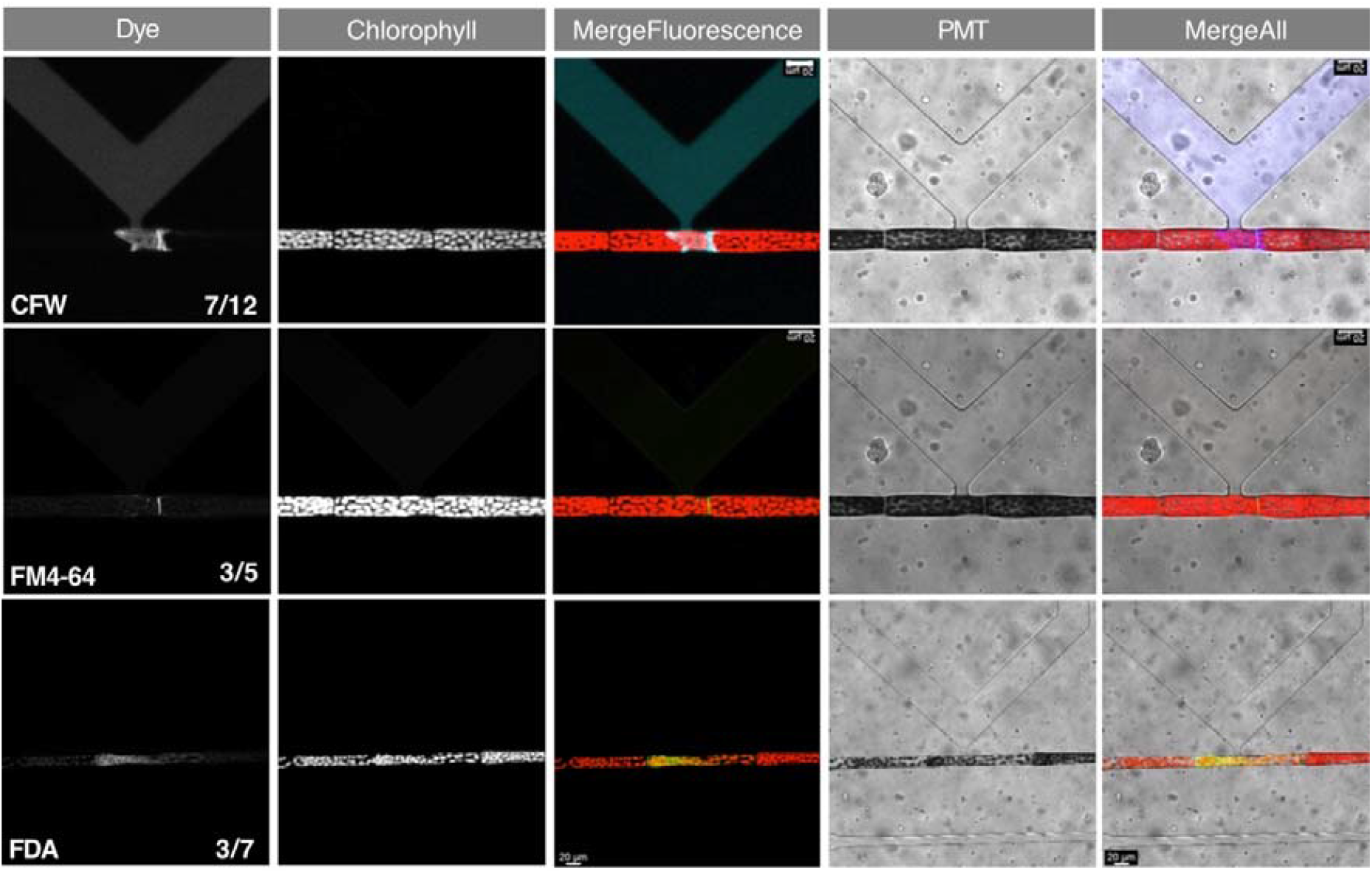
Side application of dyes, with all channels.

**Supplementary Figure 5.**
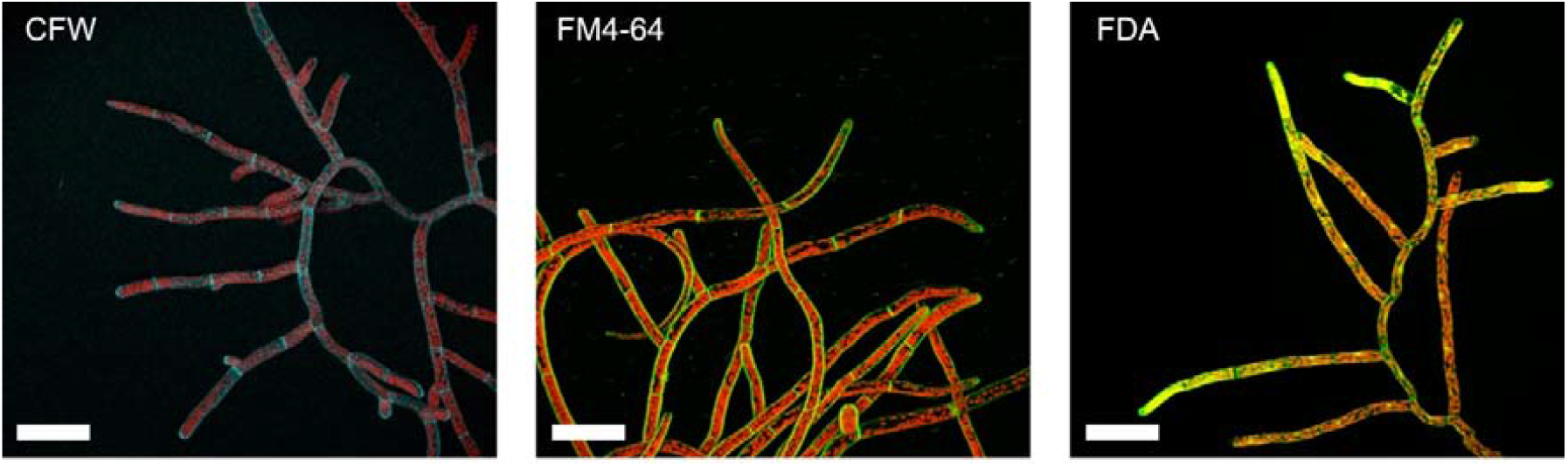
Bulk application of dyes showing variability. Calcofluor White (Left), FM4-64 (center) and Fluorescein diacetate (Right). Application of dyes in bulk shows notable variation even after repeated mixing. Comparing single cells, we see big differences that may be explained by biological variation in cell wall, membrane and metabolic activity or cell translocation, explaining why the local application of dyes does not give completely homogeneous results. High treatment delivery precision is more susceptible to biological variability.

## References

Bibeau JP, Galotto G, Wu M, Tüzel E, Vidali L. 2021. Quantitative cell biology of tip growth in moss. Plant Molecular Biology 107: 227–244.

Allan C, Sun Y, Whisson SC, Porter M, Boevink PC, Nock V, Meisrimler C-N. 2024. Observing root growth and signalling responses to stress gradients and pathogens using the bi-directional dual-flow RootChip. Lab on a Chip 24: 5360–5373.

Allan C, Tayagui A, Hornung R, Nock V, Meisrimler C-N. 2023. A dual-flow RootChip enables quantification of bi-directional calcium signaling in primary roots. Frontiers in Plant Science 13.

Ashton NW, Cove DJ. 1977. The isolation and preliminary characterisation of auxotrophic and analogue resistant mutants of the moss, Physcomitrella patens. Molecular and General Genetics MGG 154: 87–95.

Bascom CS, Wu S-Z, Nelson K, Oakey J, Bezanilla M. 2016. Long-Term Growth of Moss in Microfluidic Devices Enables Subcellular Studies in Development. Plant Physiology 172: 28–37.

Baskaran P, Soós V, Balázs E, Van Staden J. 2016. Shoot apical meristem injection: A novel and efficient method to obtain transformed cucumber plants. South African Journal of Botany 103: 210–215.

Bidhendi A j., Chebli Y, Geitmann A. 2020. Fluorescence visualization of cellulose and pectin in the primary plant cell wall. Journal of Microscopy 278: 164–181.

Brunoud G, Wells DM, Oliva M, Larrieu A, Mirabet V, Burrow AH, Beeckman T, Kepinski S, Traas J, Bennett MJ, et al. 2012. A novel sensor to map auxin response and distribution at high spatio-temporal resolution. Nature 482: 103–106.

Chen P, Li S, Guo Y, Zeng X, Liu B-F. 2020. A review on microfluidics manipulation of the extracellular chemical microenvironment and its emerging application to cell analysis. Analytica Chimica Acta 1125: 94–113.

Damme DV, Vanstraelen M, Geelen D. 2007. Cortical division zone establishment in plant cells. Trends in Plant Science 12: 458–464.

Etchells JP, Turner SR. 2010. The PXY-CLE41 receptor ligand pair defines a multifunctional pathway that controls the rate and orientation of vascular cell division. Development 137: 767–774.

Floriach-Clark J, Tang H, Willemsen V, Floriach-Clark J, Tang H, Willemsen V. 2021. Mosses: Accessible Systems for Plant Development Studies. In: Model Organisms in Plant Genetics. IntechOpen.

Grossmann G, Guo W-J, Ehrhardt DW, Frommer WB, Sit RV, Quake SR, Meier M. 2011. The RootChip: An Integrated Microfluidic Chip for Plant Science. The Plant Cell 23: 4234–4240.

Hajduk J, Szajna K, Lisowski B, Rajfur Z. 2023. The influence of microinjection parameters on cell survival and procedure efficiency. MethodsX 10: 102107.

Hiss M, Meyberg R, Westermann J, Haas FB, Schneider L, Schallenberg-Rüdinger M, Ullrich KK, Rensing SA. 2017. Sexual reproduction, sporophyte development and molecular variation in the model moss Physcomitrella patens: introducing the ecotype Reute. The Plant Journal 90: 606–620.

Holler C, Taylor RW, Schambony A, Möckl L, Sandoghdar V. 2024. A paintbrush for delivery of nanoparticles and molecules to live cells with precise spatiotemporal control. Nature Methods 21: 512–520.

Horade M, M. Kanaoka M, Kuzuya M, Higashiyama T, Kaji N. 2013. A microfluidic device for quantitative analysis of chemoattraction in plants. RSC Advances 3: 22301–22307.

Horowitz LF, Rodriguez AD, Ray T, Folch A. 2020. Microfluidics for interrogating live intact tissues. Microsystems and Nanoengineering 6: 1–27.

de Keijzer J, Freire Rios A, Willemsen V. 2021. Physcomitrium patens: A Single Model to Study Oriented Cell Divisions in 1D to 3D Patterning. International Journal of Molecular Sciences 22: 2626.

Ketelaar T. 2002. Spatial organisation of cell expansion by the cytoskeleton. Wageningen, The Netherlands.

Ketelaar T, Faivre-Moskalenko C, Esseling JJ, de Ruijter NCA, Grierson CS, Dogterom M, Emons AMC. 2002. Positioning of Nuclei in Arabidopsis Root Hairs: An Actin-Regulated Process of Tip Growth. The Plant Cell 14: 2941–2955.

Kozgunova E, Goshima G. 2019. A versatile microfluidic device for highly inclined thin illumination microscopy in the moss Physcomitrella patens. Scientific Reports 9: 1–8.

Liao C-Y, Smet W, Brunoud G, Yoshida S, Vernoux T, Weijers D. 2015. Reporters for sensitive and quantitative measurement of auxin response. Nature Methods 12: 207–210.

Mähönen AP, Tusscher K ten, Siligato R, Smetana O, Díaz-Triviño S, Salojärvi J, Wachsman G, Prasad K, Heidstra R, Scheres B. 2014. PLETHORA gradient formation mechanism separates auxin responses. Nature 515: 125–129.

Meier M, Lucchetta EM, Ismagilov RF. 2010. Chemical stimulation of the Arabidopsis thaliana root using multi-laminar flow on a microfluidic chip. Lab on a Chip 10: 2147–2153.

Menand B, Calder G, Dolan L. 2007. Both chloronemal and caulonemal cells expand by tip growth in the moss Physcomitrella patens. Journal of Experimental Botany 58: 1843–1849.

Michniewicz M, Brewer PB, Friml J. 2007. Polar Auxin Transport and Asymmetric Auxin Distribution. The Arabidopsis Book 2007.

Morejohn LC, Bureau TE, Molè-Bajer J, Bajer AS, Fosket DE. 1987. Oryzalin, a dinitroaniline herbicide, binds to plant tubulin and inhibits microtubule polymerization in vitro. Planta 172: 252–264.

Nishiyama T, Hiwatashi Y, Sakakibara K, Kato M, Hasebe M. 2000. Tagged Mutagenesis and Gene-trap in the Moss, Physcomitrella patens by Shuttle Mutagenesis. DNA Research 7: 9–17.

Remans T, Nacry P, Pervent M, Filleur S, Diatloff E, Mounier E, Tillard P, Forde BG, Gojon A. 2006. The Arabidopsis NRT1.1 transporter participates in the signaling pathway triggering root colonization of nitrate-rich patches. Proceedings of the National Academy of Sciences 103: 19206–19211.

Rensing SA, Goffinet B, Meyberg R, Wu S-ZZ, Bezanilla M. 2020. The moss physcomitrium (Physcomitrella) patens: A model organism for non-seed plants. Plant Cell 32: 1361–1376.

Scott AC, Allen NS. 1999. Changes in Cytosolic pH within Arabidopsis Root Columella Cells Play a Key Role in the Early Signaling Pathway for Root Gravitropism1. Plant Physiology 121: 1291–1298.

Smith LG. 2001. Plant cell division: building walls in the right places. Nature Reviews Molecular Cell Biology 2: 33–39.

Stanley CE, Shrivastava J, Brugman R, Heinzelmann E, van Swaay D, Grossmann G. 2018. Dual-flow-RootChip reveals local adaptations of roots towards environmental asymmetry at the physiological and genetic levels. New Phytologist 217: 1357–1369.

Thelander M, Landberg K, Sundberg E. 2019. Minimal auxin sensing levels in vegetative moss stem cells revealed by a ratiometric reporter. New Phytologist 224: 775–788.

Wang Y, Dindas J, Rienmüller F, Krebs M, Waadt R, Schumacher K, Wu W-H, Hedrich R, Roelfsema MRG. 2015. Cytosolic Ca2+ Signals Enhance the Vacuolar Ion Conductivity of Bulging Arabidopsis Root Hair Cells. Molecular Plant 8: 1665–1674.

Wang JD, Douville NJ, Takayama S, ElSayed M. 2012. Quantitative Analysis of Molecular Absorption into PDMS Microfluidic Channels. Annals of Biomedical Engineering 40: 1862–1873.

Yanagisawa N, Higashiyama T. 2018. Quantitative assessment of chemotropism in pollen tubes using microslit channel filters. Biomicrofluidics 12: 024113.

Yanagisawa N, Sugimoto N, Arata H, Higashiyama T, Sato Y. 2017. Capability of tip-growing plant cells to penetrate into extremely narrow gaps. Scientific Reports 7: 1403.

Yoshida MW, Kozgunova E. 2023. Microfluidic Device for High-Resolution Cytoskeleton Imaging and Washout Assays in Physcomitrium (Physcomitrella) patens. In: Hussey PJ, Wang P, eds. Methods in Molecular Biology. The Plant Cytoskeleton: Methods and Protocols. New York, NY: Springer US, 143–158.

